# Accurate analysis of short read sequencing in complex genomes: A case study using QTL-seq to target blanchability in peanut (*Arachis hypogaea*)

**DOI:** 10.1101/2021.03.13.435236

**Authors:** Walid Korani, Dan O’Connor, Ye Chu, Carolina Chavarro, Carolina Ballen, Peggy Ozias-Akins, Graeme Wright, Josh Clevenger

## Abstract

Next Generation sequencing was a step change for molecular genetics and genomics. Illumina sequencing in particular still provides substantial value to animal and plant genomics. A simple yet powerful technique, referred to as QTL sequencing (QTL-seq) is susceptible to high levels of noise due to ambiguity of alignment of short reads in complex regions of the genome. This noise is particularly high when working with polyploid and/or outcrossing crop species, which impairs the efficacy of QTL-seq in identifying functional variation. By filtering loci based on the optimal alignment of short reads, we have developed a pipeline, named Khufu, that substantially improves the accuracy of QTL-seq analysis in complex genomes, allowing *de novo* variant discovery directly from bulk sequence. We first demonstrate the pipeline by identifying and validating loci contributing to blanching percentage in peanut using lines from multiple related populations. Using other published datasets in peanut, *Brassica rapa, Hordeum volgare, Lactua satvia*, and *Felis catus*, we demonstrate that Khufu produces more accurate results straight from bulk sequence. Khufu works across species, genome ploidy level, and data types. In cases where identified QTL were fine mapped, the fine mapped region corresponds to the top of the peak identified by Khufu. The accuracy of Khufu allows the analysis of population sequencing at very low coverage (<3x), greatly decreasing the amount of sequence needed to genotype even the most complex genomes.

## Introduction

Quantitative Trait Locus Sequencing (QTL-seq) is a genetic analysis that takes advantage of the ever-decreasing cost in whole genome sequencing (WGS). The two seminal papers describing the simple approach of bulk segregant analysis were first published in 1991 (Giovannoni *et al.*, 1991; Michelmore *et al.*, 1991). The markers being used were restriction fragment length polymorphism (RFLP) or random amplified polymorphic DNA (RAPD), and in the absence of genome sequences, identifying new markers was difficult within mapping populations. After an interval was initially mapped to control a trait of interest, it was not possible to fine map the interval further in the absence of polymorphic markers. The method of bulk segregant analysis (BSA) rested on the hypothesis that markers linked to the trait of interest will segregate with the phenotype in a segregating population. The individuals exhibiting the extreme tails of the phenotypic distribution will be differentiated by only those markers that were physically within the genomic region of interest. In the case of the original methods, random oligos could be screened in the bulked DNA until polymorphic markers were found. This was how new markers could be identified and placed on the genetic map to fine map the trait to a higher resolution. The application of BSA evolved as molecular markers became higher throughput. Hybridization expression arrays were used in *Arabidopsis* to map mutations (Borevitz *et al.*, 2003). In yeast, hybridization arrays and the ability to assay extremely large populations allowed profiling the structure of complex traits controlled by many loci (Ehrenreich *et al.*, 2010). Whole genome sequencing (WGS) using short reads resulted in more accuracy for mapping mutations in *Arabidopsis* (Ossowski *et al*, 2008; Schneeberger *et al.*, 2009). This was extended to plants without reference genomes as mapping-by-sequencing (Galvão *et al.*, 2012). Mapping mutations in crop plants with larger genomes was introduced as MutMap, which highlighted rice (Abe *et al.*, 2012), and extended using mutants without crossing back to a wild type parent (Fekih *et al.*, 2013). Using rice, BSA was used in combination with WGS to map natural variation in crops in a technique named QTL-seq (Takagi *et al.*, 2013). The application of QTL-seq moved to more complex genomes; tomato (Illa-Berenguer *et al.*, 2015), chickpea (Das *et al.*, 2015), cucumber (Lu *et al.*, 2014), oilseed rape (Wang *et al.*, 2016), guinea yam (Tamiru *et al.*, 2016), watermelon (Gimode *et al.*, 2020), and peanut (Pandey *et* al., 2017; Clevenger *et al.*, 2018; Kumar *et al.*, 2020; Zhao *et al.*, 2020).

The strengths of QTL-seq are its simplicity, efficiency of resource use, and flexibility to utilize historical data. There are limitations however. If the parental genotypes used are divergent from the reference genome, there is variation that short reads cannot account for. This variation; insertions, deletions, duplications, and copy number variants, has major consequences for the expression of phenotypes (Schiessl *et* al., 2019; Alonge *et al.*, 2020). If one parent is more divergent from the reference than the other parent, then reference bias can skew allele frequencies away from the true frequency in the bulks (Günther and Nettelblad, 2019). For crop genomes, which are either polyploid or have a polyploidization event in their evolutionary history, homeologous sequences introduce an additional layer of noise (Glover *et al.*, 2016; Clevenger *et al.*, 2015). The consequence is that short reads will pile up in regions that are duplicated in the bulks, but not in the reference, resulting in incorrect allele frequency for that physical location. Alternatively, homeologous sequences will map incorrectly and create further noise in the signal.

For complex polyploid genomes like peanut (*Arachis hypogaea*), the analysis of the bulk sequence data has relied on generating high quality parental sequence data first, and then utilizing it to either modify the reference genome to be parental-specific (Pandey *et* al., 2017; Kumar *et al.*, 2020; Zhao *et al.*, 2020), or to call SNPs between the parents with specialized pipelines to use in the analysis of the bulks (Clevenger *et al.*, 2018; Cui *et al.*, 2020). In heterozygous crops, like *Brassica rapa*, a strategy was devised that used sequence from parents and F1 hybrids to identify SNPs that were homozygous in the parent of interest (Itoh *et al.*, 2019). In the case of peanut, the results have been useful, but still exhibit noise to the extent that the precise structure of the identified loci are not clear. The higher noise to signal ratio also precludes identifying minor effect QTL. In cases where the parents of a population are ambiguous, or when using breeding populations with complex pedigrees, it is more useful to be able to call SNPs directly from the bulk sequence without needing parental sequence. Relying on parental sequence is not cost effective, and it also limits the genetic material that can be used to map traits. For complex genomes, it is imperative to be able to analyze QTL-seq datasets from the bulk sequence alone.

A useful outcome for QTL-seq analyses is that markers can be developed straight from breeding populations to be used directly within those breeding programs (Pandey *et al.*, 2017; Clevenger *et al.*, 2018; Cui *et al.*, 2020). A good example of this is rust resistance in the breeding programs at ICRISAT in India (Pandey *et al.*, 2017). The markers developed from QTL-seq analysis have been included in a SNP panel that is being used in the global peanut breeding community to select for rust resistance. In this case, it was the discovery of a wild introgression that conferred resistance and was amenable to marker-assisted selection with markers that could select for the introgression on chromosome A03 (Pandey *et al.*, 2017). A trait that is important for peanut processing and is time consuming to measure is blanchability (Wright *et al.*, 2018; Cruickshank *et al.*, 2003). Blanching consists of heating peanuts to a temperature that removes the skins without damaging the kernel. It is very important to blanch peanuts for quality control (removal of damaged kernels, including aflatoxin contamination) and for processing applications (peanut butter, confectionery). Wright and colleagues (2018) found that the blanching percentage was under strong genetic control with very low genotype x environment (G x E) interactions. Peanut breeding programs generally do not select for blanching percentage due to the high labor investment needed to test. Additionally, a large number of seeds are needed to test blanching percentage, which precludes selection at early generations. Current uniform trials in the United States do not test for blanching percentage (https://www.ars.usda.gov/southeast-area/dawson-ga/national-peanut-research-laboratory/docs/uniform-peanut-performance-tests-uppt/). Despite not having information on the blanching percentage of different cultivars, the cost savings for manufacturers would be substantial. After blanching, any unblanched kernels are discarded as waste, sold as less valuable products, or crushed for oil. The combination of strong genetic control, small G x E effect, and laborious measurement makes blanching an excellent target for marker assisted selection (MAS) in peanut (Wright *et al.*, 2018; Cruickshank *et al.*, 2003). In this study, we report a method for filtering SNPs effectively to reduce noise in the analysis of QTL-seq experiments. The method identifies high quality SNPs by concentrating on alignment accuracy and allele accuracy at each locus. Each *de novo* identified SNP is then profiled directly from bulk sequence to calculate the allele frequencies. We demonstrate the pipeline by re-analyzing published peanut datasets without using parental sequence and show increased accuracy. We identify previously unidentified minor QTL and validate them. We use the pipeline to *de novo* analyze a new dataset focused on blanching in peanut and validate those results in an independent breeding population. We analyze published datasets from outcrossing crop species, inbred species with large genomes, and domestic cats. These datasets use different sequencing strategies, including WGS, exome capture, and RNA sequencing. We further show that our pipeline can analyze low coverage sequencing effectively using examples in peanut. The pipeline, named after the Khufu pyramid, is capable of *de novo* analysis of QTL-seq datasets using complex genomes, unlocking the potential of QTL-seq in complex populations without the need for additional sequencing.

## Results and Discussion

### Blanchability QTL-seq

To identify QTLs associated with blanchability in peanut, we sequenced two bulks of breeding lines from three populations with related parents (Table 1). The parents with high blanching percentage were Walter and Redvale and the low blanching parents were Sutherland and a sister line of Sutherland (Wright *et al.*, 2018). Redvale is a selection from a cross with Walter as a parent. We first analyzed the data by calling parent-specific SNPs by re-sequencing Walter, Redvale, and Sutherland and identifying SNPs where Walter and Redvale shared an allele and Sutherland had an alternate allele. We used these SNPs to analyze the QTL-seq dataset and identified two QTLs on chromosomes A06 and B11 (Figure 1A). The parents of the population are Sutherland (poor blancher; 74.5%) and Middleton (good blancher; 89%). Middleton is related to Walter and Redvale, but not a sister line. Markers were developed from the most significant SNPs within the identified peaks on A06 and B01. Both markers could select for increased blanching percentage in an unrelated RIL population, with A06 having a stronger effect (F=6.73; df=176; RANK ANOVA p=0.0002; Kruskal p=0.0003; Effect 9.8%; Median=73.4%; R^2^=10.3%) than B01 (F=3.5; df=176; RANK ANOVA p=0.01; Kruskal p=0.005; Effect 8.03%; Median=74%; R^2^=5.6%). The two markers combined were additive (F=5; df=176; RANK ANOVA p<0.0001; Effect 19.7%; Median=76.9%; R^2^=14.8%). The lsmean estimate when both alleles are beneficial is 79.3% and when both alleles are non-beneficial is 59.6%. Selecting with the two markers shifts the median blanching percentage from 62.8% to 83.9% (21% increase).

**Table 1:** Phenotypes for selected lines for bulk sequencing.

| **Genotype** | **Blanching** | **Pedigree** | **Bulk** |
| --- | --- | --- | --- |
| P24-p188-93 | 55.4 | Tingoora x D147-p3-115 | Poor Blancher |
| P13-p07-219 | 59.6 | Sutherland x Walter | Poor Blancher |
| P24-p187/205-97 | 71.5 | Tingoora x D147-p3-115 | Poor Blancher |
| P23-p157-65 | 74.1 | Redvale x D147-p3-115 | Poor Blancher |
| P13-p45-235 | 76.7 | Sutherland x Walter | Poor Blancher |
| P23-p157-64 | 84.0 | Redvale x D147-p3-115 | Good Blancher |
| P13-p23-233 | 85.2 | Sutherland x Walter | Good Blancher |
| P23-p153-62 | 85.8 | Redvale x D147-p3-115 | Good Blancher |
| P13-p21-223 | 91.0 | Sutherland x Walter | Good Blancher |
| P23-p171-85 | 91.0 | Redvale x D147-p3-115 | Good Blancher |
| Walter | 92.2 | Walter x D45-p37-102 | Good Blanching Parent |
| Redvale | 93.3 |  | Good Blanching Parent |
| Sutherland* | 75.4 |  | Poor Blanching Parent |

\* Data from 2019

**Figure 1:**
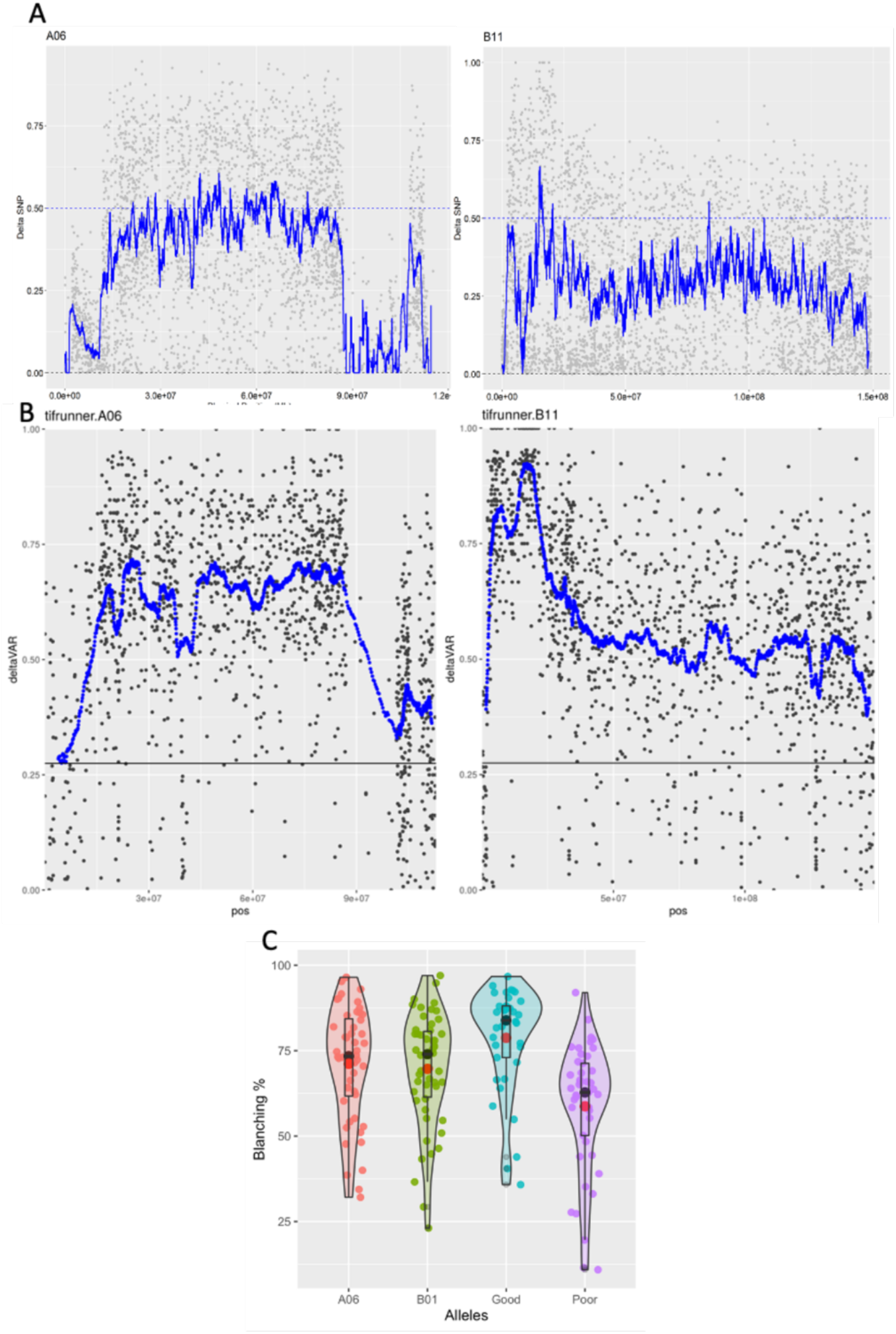
**A)** Initial QTL-seq using parental sequence to identify SNPs for use in analyzing bulk sequence. Delta SNP is 0 to 1 where 1 is high blanching allele. **B)** The same QTLs identified with Khufu *de novo* from bulk sequence. Because allele parentage is not discerned, delta SNP is set from 0 (equal frequency between bulks) to 1 (complete segregation between bulks). **C)** Validation of markers in an independent RIL population. Violin/Jitter/Box plots show the blanching % of RILs with the high blanching allele at A06 only, B01 only, Both A06 and B01, and both poor (low blanching) alleles. Black dots show median values and red dots are the mean. Selecting for either marker alone is significant compared to selecting for both poor alleles and selecting for the high blanching allele at both loci provides the highest blanching percentage and skews the distribution towards 100%.

We used Khufu to re-analyze the blanching QTL-seq dataset using the *de novo* discovered SNPs from the bulk data (Figure 1B). The QTL on A06 has a stronger signal overall with the smoothed average above 0.6 with Khufu compared to 0.5 using parental sequence. The QTL on B11 is much stronger with the peak smoothed average reaching 0.85 compared to 0.65 using parental data. The peak on B01 is also clearly defined using Khufu.

### Peanut stem rot QTL-seq

Cui *et al.* (2020) used QTL-seq to identify QTLs contributing to stem rot resistance in the field in peanut. There were two QTLs identified on chromosomes A05 and A01 that could explain about 21% of the variance observed. The strategy used in Cui *et al.* (2020) was to identify putative parental SNPs from parental sequence (using the polyploid-specific pipeline SWEEP (Clevenger *et al.*, 2018)) that were then used in the bulks to calculate allele frequency. Using Khufu, we analyzed the bulk sequence *de novo* and identified the peaks on chromosomes A01 and A05 (Figure S1). On chromosome A01, the smoothed average within the peak was consistently above 0.6 in contrast to the original analysis where the smoothed average was a jagged line that was above 0.5 at several positions, but not consistently. There is an additional peak on the bottom of the chromosome outside of the pericentromeric region that is clearly significant in the Khufu analysis where noise washed it out in the original analysis. Khufu identified two additional peaks on chromosomes A04 and A07 that were not identified previously (Figure S2). We developed Kompetitive Allele Specific PCR (KASP) markers to assay the new QTL on A04 in the validation populations used in Cui *et al* (2020). These markers were able to increase the ability to select for stem rot resistance (Figure S2).

### Additional examples of improved QTL resolution

The stem rot bulks were sequenced to high depth (>40X genome coverage). We re-analyzed a QTL-seq dataset focused on late leaf spot resistance in peanut where the bulks were sequenced to approximately 10X genome coverage (Clevenger *et al.*, 2018). Khufu successfully identified the three QTL *de novo* from bulk sequence that were validated in Clevenger *et al.* (2018) and Chu *et al.* (2019) (Figure S3). The QTL on B03 (13) has a clear peak in the Khufu analysis indicating potential to develop more tightly linked markers to resistance. Pandey *et al* (2020) used QTL-seq to identify QTLs associated with fresh seed dormancy in peanut. Using Khufu, we identified those same QTLs *de novo* from the bulked sequence (Figure S4). As was the case with the stem rot dataset, the Khufu peaks had a higher smoothed average on chromosome B05, reaching 0.75 in the region from which Pandey and colleagues (2020) developed a marker to select for the trait. Zhao and colleagues (2020) used QTL-seq to identify a candidate gene for purple testa color in peanut on chromosome A10 using parental sequence to analyze the bulk sequence data. Without parental sequence, Khufu identified the same QTL and the predicted peak region (107,280,934 to 110,087,506) contains the most likely candidate gene which was functionally shown to control anthocyanin accumulation in tobacco (Zhao *et al.*, 2020).

Khufu is effective in other crops with complex genomes other than peanut. We re-analyzed the data from Itoh *et al.* (2019) where they reported a strategy to analyze QTL-seq data in *Brassica rapa*, a highly heterozygous species. They identified parental SNPs by first sequencing the parents of the population, and filtered those SNPs by identifying those that were heterozygous in the F1 hybrid using sequence of the F1. Using Khufu, we identified the same two QTL on chromosomes A07 and A10 straight from the bulk sequence with no need for parental or hybrid sequence (Figure S6).

We investigated using different data types that reduce the complexity of the genome. Two groups used exome capture to map QTL in Barley (Hisano *et al.*, 2018; Kodama *et al.*, 2018). Hisano *et al* (2018) looked at net blotch resistance and the black kernel trait (*blp*). They used a concatenated transcriptome organized by physical distance. We analyzed those datasets using the *Hordeum vulgare* reference genome sequence. For net blotch, there are two QTL located in pericentromeres of chromosomes 3 and 6 (Figure S7). For *blp*, Long and colleagues (2019)fine mapped the mutation to a region including 537,855,051 to 538,661,850. Using the data from Hisano *et al* (2018), Khufu predicts the region to be between 535,000,000 and 540,000,000, which contains the fine mapped region identified by Long *et al* (2019) (Figure S8). The tip of the peak indicated by Khufu is nearest a SNP within a candidate gene identified by Long and colleagues (1H:538,502,975, HORVU1Hr1G087010). Kodama and colleagues investigated fertility under salt stress (Figure S9). Khufu analysis presented a clearer picture despite the two QTL mapping to the pericentromere. Finally, Khufu can process RNA-seq data effectively, highlighted by mapping genes controlling red leaf color in lettuce (*Lactuca sativa*; Figure S10). Khufu also is effective for mapping traits in domestic cats. Graff *et al* (2020) identified a mutation in *PEA15* that leads to impaired cerebral cortical size in domestic cats by analyzing sequence data from a set of affected and unaffected individuals. We analyzed only the WGS data which included 3 obligate carriers and 2 affected cats and RNA-seq data including 7 affected cats and 4 unaffected cats (Figure S11). Khufu successfully identified the region on chromosome F1 including *PEA15*.

Khufu is successful at analyzing bulk sequencing datasets in complex genomes without needing additional sequence (parents, F1 hybrid, etc), which streamlines QTL-seq analysis, makes it more powerful, and opens up new populations with complex parentage (multi parent populations) and breeding populations where parental genotypes are less certain. Bertioli *et al.* (2020) estimated that the rate of polymorphism between two peanut accessions is on average 1 SNP per 10,000 bp by comparing two reference-quality genome sequences. One advantageous aspect of QTL-seq is the theoretical ability to have access to all SNP polymorphisms segregating in the population. The result is that the SNP or group of SNPs that are most closely linked to the functional variation (or are the functional variation) are available. In contrast, the probability that a SNP array with fixed markers includes the most closely linked SNPs is low and decreases with low throughput marker types. Khufu identified 244,329 SNPs in the dormancy dataset (1 SNP per 10,395 bp), 372,611 SNPs in the stem rot dataset (1 SNP per 6,816 bp), and 225,029 SNPs in the purple testa dataset (1 SNP per 11,287 bp). The number of SNPs Khufu filtering identifies *de novo* from the bulk sequences is within the estimate of polymorphic SNPs between any two *A. hypogaea* accessions. The filtering procedure of Khufu should be applicable for other genotyping applications using short reads in crops with complex genomes.

### Khufu filtering compared to map quality-based filtering

Short read aligners such as BWA (Li and Durbin, 2009) calculate a quality score based on the confidence that a read maps to a location in the genome. The quality is reported as the phred-based probability that the read is mapped incorrectly. A strategy for ensuring uniquely mapped, highly confident mapping is to filter using map quality. We will refer to this as the “standard approach” (Figure 2). Compared to Khufu filtering using the blanching bulk sequence, the standard approach includes substantial noise in the peak region (false positive SNPs with dSNP values < 0.3 within the peak; Figure 2). Khufu filtering shifted the signal to noise ratio from 1:4 with the standard approach to greater than 4:1 with Khufu (Figure S12), indicating a substantial increase in accuracy.

**Figure 2:**
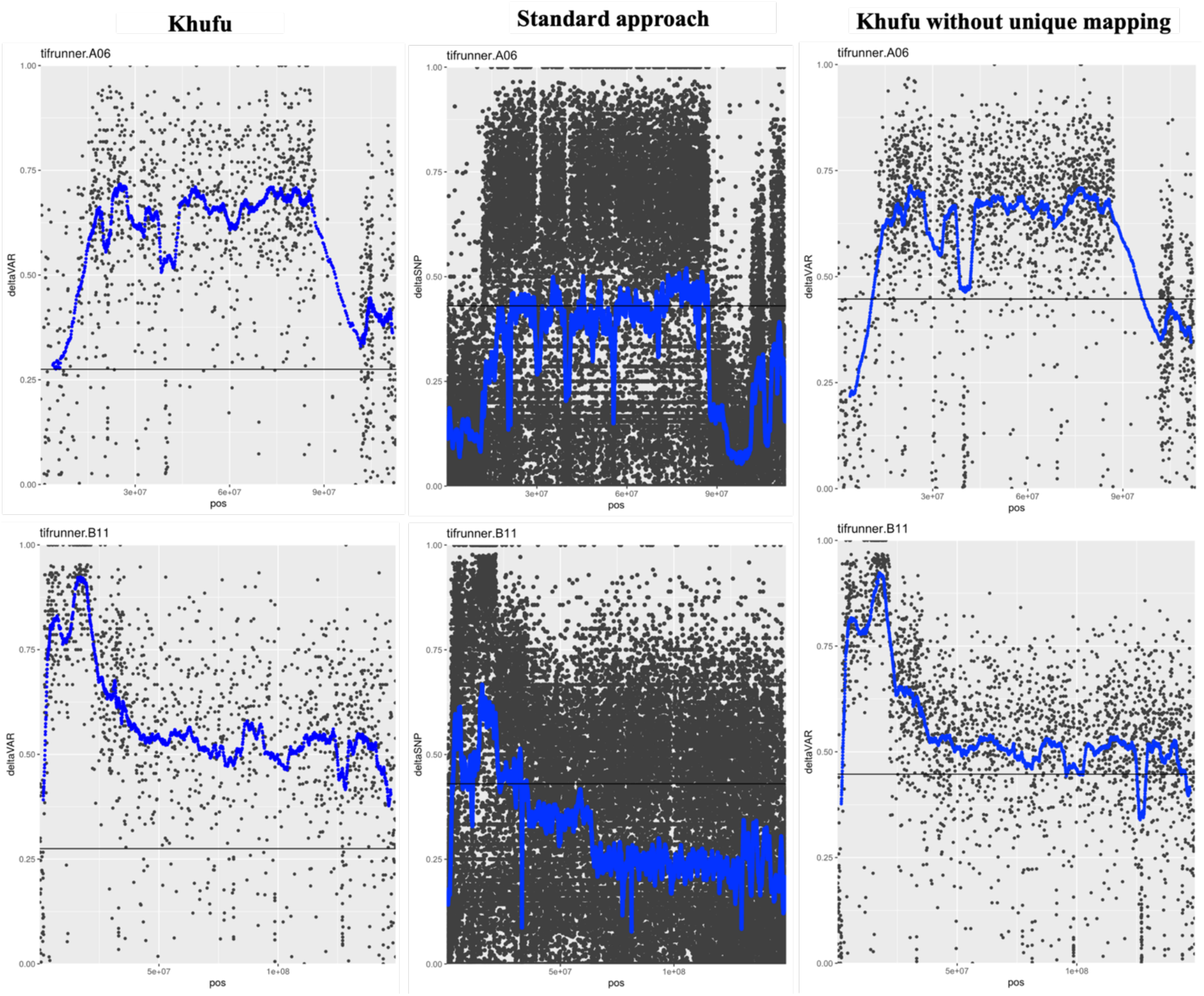
Dissecting the importance of each step in the pipeline. Full dataset refers to running the complete pipeline on the blanching data. Shown are the two QTLs on A06 and B11. Standard Approach refers to filtering using map quality and unique mapping. Khufu includes filtering for unique mapping. Khufu without unique mapping is the Khufu pipeline without filtering for uniquely mapping reads first.

We next asked the question, how much effect does filtering for map quality and constraining reads to unique mapping have on Khufu (Figure 2)? Filtering for uniquely mapped reads has little to no effect on the final analysis, showing the filtering strategy of Khufu is not successful due to only keeping reads with high map qualities.

### Considerations of sequencing depth

The goal of QTL-seq is to assess the allele frequency of the individuals comprising each bulk. At least one sequenced read per individual would be required to estimate the allele frequency accurately. Even though sequencing cost has decreased substantially, many researchers still try to weigh the tradeoffs between higher sequencing depth and cost savings. Using the blanching dataset, which was sequenced to a common minimum depth of 31X coverage, fidelity is retained as low as 20X coverage (Figure 3). At 10X coverage, the peaks are still significant, although the smoothed average is about 10% lower, which will have an impact on the ability to identify minor effect loci. We re-analyzed published peanut datasets that were sequenced to 10X coverage and attained good results (Figures S3, S4). Sequencing as low as 10X coverage will produce acceptable results, however the lowest depth that produces a clear result is 20x coverage. This result is the same for the blanching dataset where 5 individuals were pooled (4 reads per individual) and the stem rot dataset where 10 individuals were pooled (2 reads per individual). Even at 5X coverage, Khufu successfully identifies the significant QTLs, albeit at lower fidelity. At such low depth it is more a lack of accuracy in allele frequency, where there are only 6 possible frequencies possible with only 5 reads available per site.

**Figure 3:**
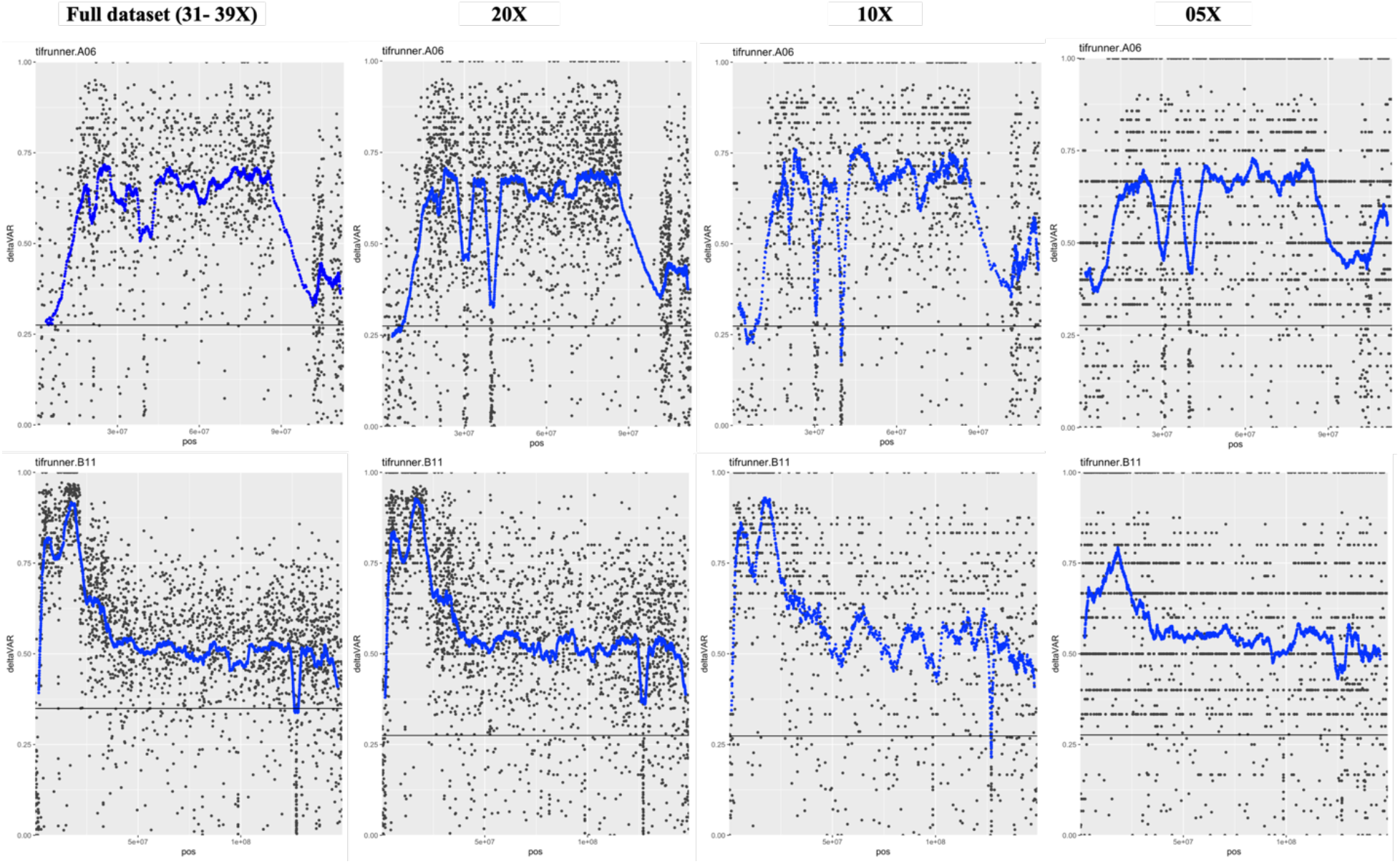
How depth affects peak resolution. The full blanching % dataset (31 – 39x coverage) was down sampled to 20X, 10X, and 5X. Shown are QTl on A06 (top) and B11 (bottom).

### Potential for genotyping using skim sequencing

For crops with large, complex genomes, genotyping large populations still relies heavily on fixed SNP arrays. These arrays are incredibly useful, yet suffer from ascertainment bias. Reduced representation methods allow *de novo* SNP discovery yet only access a small portion of the genome. The accuracy of profiling SNPs within bulked samples should be useful for genotyping applications using sequencing. As a reference set of SNPs, we identified variants by comparing two published peanut genomes, Tifrunner (Bertioli *et al.*, 2019), and Shitouqi (Zhuang *et al.*, 2019). Using Illumina sequencing of the two genotypes that were sequenced, we called SNPs using Khufu at different depths of coverage (Table S1). Across all levels of depth, Khufu calls alleles with more than 99% accuracy (according to the allele in the reference genome sequence). Above 10X coverage, Khufu identifies more than 95% of the SNPs. At lower coverage, Khufu identifies more than 71% at 5X coverage and 46% at 3X.

We tested if it were possible to identify *de novo* SNPs in the parents of a population sequenced at 10X coverage and then call alleles in the progeny sequenced at 1X or even 0.5X coverage (Table S2). At 1X coverage, 27% of the possible SNPs could be called at 100% allele accuracy. At 0.5X coverage, 14% of possible SNPs could be called at 100% allele accuracy. With an estimate of 1 SNP per 10 kilobases polymorphic between any two peanut accessions, we can expect to call an average 67,000 SNPs with 1X coverage and 37,500 with 0.5X coverage. This represents genome-wide distribution of markers.

Finally, we analyzed a RIL population that was genotyped using WGS re-sequencing (Agarwal *et al.*, 2019). Using 5x coverage WGS of the progeny, Agarwal and colleagues mapped 11,106 SNPs using SWEEP_GOLD (https://github.com/w-korani/SWEEP_GOLD). Using Khufu, we identified and called 86,987 SNPs in the progeny after down sampling to <3x coverage.

Analysis of physical maps color-coded for parental allele illustrates the accuracy of Khufu at low coverages (Figure S13).

### Conclusions

Profiling SNP variants accurately straight from bulk sequence data is critical for analyzing QTL-seq experiments. The efficiency of QTL-seq lies in the simplicity of sequencing only two samples. It becomes more complicated and expensive when additional samples need to be sequenced. We present a method to accurately analyze QTL-seq datasets by identifying SNPs *de novo* straight from bulk sequences. We show by re-analyzing public datasets of crops with complex genomes that the presented method is more accurate using only bulk sequence than sequencing parent lines. Further, we demonstrate that the presented method is superior to filtering procedures available from mapping scores. This method, which we call Khufu, named after the Great Pyramid, has an additional application to accurately genotype lines with skim sequencing. Khufu truly unlocks the power of QTL-seq to efficiently identify QTL associated with traits of interest.

## Materials and Methods

### Post-Khufu filtering

We implemented a shiny app to facilitate final polishing post Khufu processing. It can be freely accessed here: https://w-korani.shinyapps.io/khufu_var2/. This allows filtering in real time so the user can identify trends within potential noisy groups of SNPs. There is minimum and maximum depth filter. Low depth loci estimate allele frequencies poorly and loci with depths more than twice the estimated read coverage can signal regions of paralogous read mapping. The minor allele frequency filter was first suggested by Takagi *et al* (2013) as a mechanism to filter erroneous base calls if a particular alternative allele was in low frequency in both bulks. We implemented an “interval” filter which filters out potential SNPs clustered closely together (<100 bp). This indicates repetitive content and reads from many loci overlapping one locus. The shiny app takes as input R objects that are created by Khufu. In the supplement we have included the R objects for the full Blanching data set, and the down sampled 5X, 10X, and 20X sets.

### Peak Confidence Intervals

The shiny app includes an “interactive peak” option. During the analysis of published datasets, it became clear that the top of the peaks corresponded to where regions were ultimately fine mapped following up the initial QTL-seq analysis. We devised a simple test to assign confidence intervals to a selected peak region. We first tested windows of dSNP values genome wide to determine if they followed a normal distribution using a Shapiro-Wilks test. The average p-value genome wide was very significant so we implemented a non-parametric test to test for significance between two samples of dSNP distributions. The user picks the top of the peak and the test moves to the left and right, testing if the set of dSNP values is significantly different from the one before using a Kruskal Wallis test. The confidence intervals are set at the locations where a significant difference is found, indicating that the following set of allele frequencies are significantly lower (less linked to the phenotype) than the peak.

### Blanching QTL-seq and validation

Good and poor blanching lines were selected from data generated in Wright *et al.*, 2018. Each line was extracted for DNA individually and quantified using a Qubit fluorometer. Equal concentrations of each were pooled into ‘good blanching’ and ‘poor blanching’ DNA pools. The pools were sequenced by Novogene using Illumina HiSeq 2×150 bp sequencing. After filtering, a total of 193,712,311 reads from the good blanching pool were mapped (100%; 94.9% properly paired) and 248,926,026 reads from the poor blanching pool (100%; 94.9% properly paired). Analysis of sequenced bulks was carried out using the Khufu pipeline.

To compare Khufu to a traditional approach, good and poor blanching pools were mapped to the Tifrunner genome v2 (Bertioli *et al.*, 2019; peanutbase.org) using BWA (Li and Durbin, 2009) and mapped reads were filtered for unique alignment (grep -v ‘XA:Z:’|grep -v ‘SA:Z:’ |awk ‘{if($5==60 || $5 == “”) print($0)}’) where $5 represents the map quality (60 is highest quality BWA assigns). Further depth filtering was at least 5 reads covering the SNP and a maximum of 2 times the number of reads covering the SNP compared to the expected read depth. Khufu was run in full without filtering for unique reads to compare the impact of unique mapping to the Khufu pipeline.

### Development of RIL population and marker validation

A recombinant inbred line (RIL) population was developed by crossing Middleton x Sutherland cultivars, comprising of 406 individual lines (RILs) at the Peanut Company of Australia (PCA), Kingaroy, Queensland, Australia. Middleton is a reliable high yielding commercial cultivar and has been used extensively in the Australian Peanut Breeding Program (APBP). The pedigree of the resistant parent, Sutherland, is (B123 (F1) × CS22) × D45-p37-102, and is tolerant to multiple foliar diseases; late leaf spot (*Phaeoisariopsis personata*), peanut rust (*Puccinia arachidis*) and Web Blotch (Khedikar et al. 2010). Hybridization between parental lines was performed in 2014 and the single seed descent (SSD) method was employed under field conditions from the F2 to the F6 generation. Extracted DNA from 406 RIL lines were used for marker validation. Blanching percentage was performed as described in Wright *et al.*, 2018. Kompetitive Allele Specific PCR (KASP) markers were designed from the most significant SNPs from the QTLs on B01 and A06 (Table S1). Two methods were used to assess the significance of each identified locus to blanching percentage. The blanching data were transformed into ranks and ANOVA’s were run using each marker alone and the combination of both markers in R. LSMeans for each marker and combinations of them were calculated using the lsmeans() package in R. An additional test using a Kruskal Wallis non parametric test was performed using Kruskal.test() in R.

## Supporting information

Supplemental_khufu_figures&tables

blanching_R_objects_4sets

## Analysis of published datasets

Sequences from published datasets were downloaded from public databases (NCBI SRA and DDBJ Bioproject). Raw fastq files were not processed further and were subjected to the Khufu pipeline as downloaded.

## Data availability

R object files of blanching data of the full set, and 20X, 10X, and 5X subsets are included in the supplemental materials. Raw data fastq sequences of high and low bulks of blanching data are deposited at the NCBI (http://www.ncbi.nlm.nih.gov/) under BioProject PRJNA714121.

## Funding

Funding was provided by Grains Research and Development Corporation-GRDC, under project ‘Australian Peanut Breeding Program - PCA00003.

## Acknowledgments

We acknowledge Justin N. Vaughn for discussing the methods used and reviewing the manuscript, and Mars-Wrigley for supporting J Clevenger for a portion of the project. We also acknowledge Professor Robert Henry, Dr Agnello Furtado and Dr R.C.N. Rachaputi from the Queensland Alliance for Agriculture and Food Innovation (QAAFI) at the University of Queensland, Australia, for their preliminary work on development of blanching markers.

## Author Contribution

WK and JC analyzed published datasets, developed Khufu, and drafted the manuscript. WK designed and wrote the code for the shiny application and for all modules of Khufu. YC and POA validated QTL and revised the manuscript. CC and CB provided technical lab support. DOC conceived and carried out the blanching experiments. JC and GW conceived and supervised the project, secured funding, and revised the manuscript.

## Notes

### Competing Interest Statement

The authors have declared no competing interest.

