## Supplemental_khufu_figures&tables for "Accurate analysis of short read sequencing in complex genomes: A case study using QTL-seq to target blanchability in peanut (*Arachis hypogaea*)"

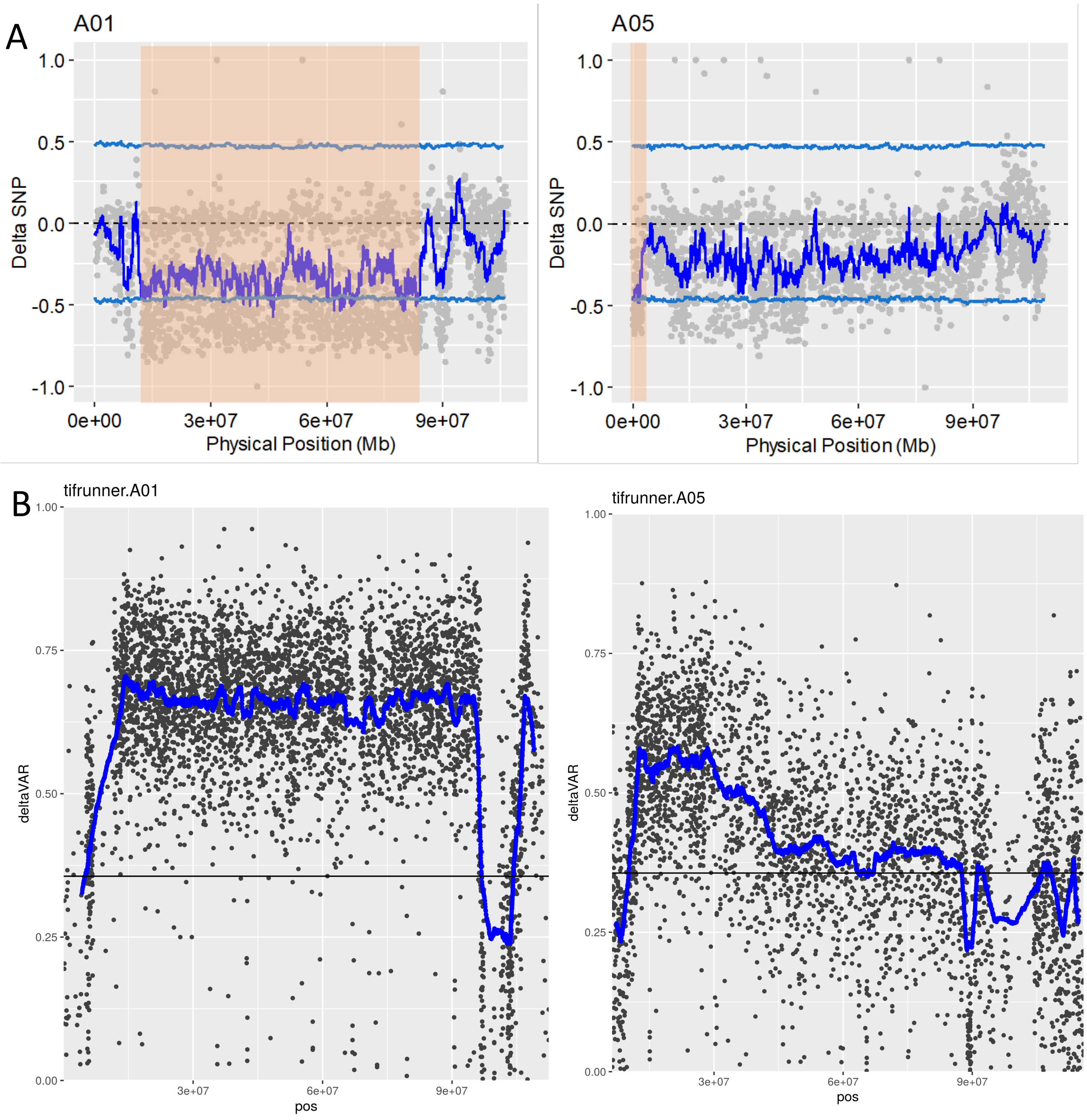


**Figure S1: A)** Original dSNP plots from Rui *et al.*, 2019 where SNPs were identified from parental sequence first and then assayed in the bulks. **B)** dSNP plots from khufu showing more clear identification of QTLs without parental sequence.

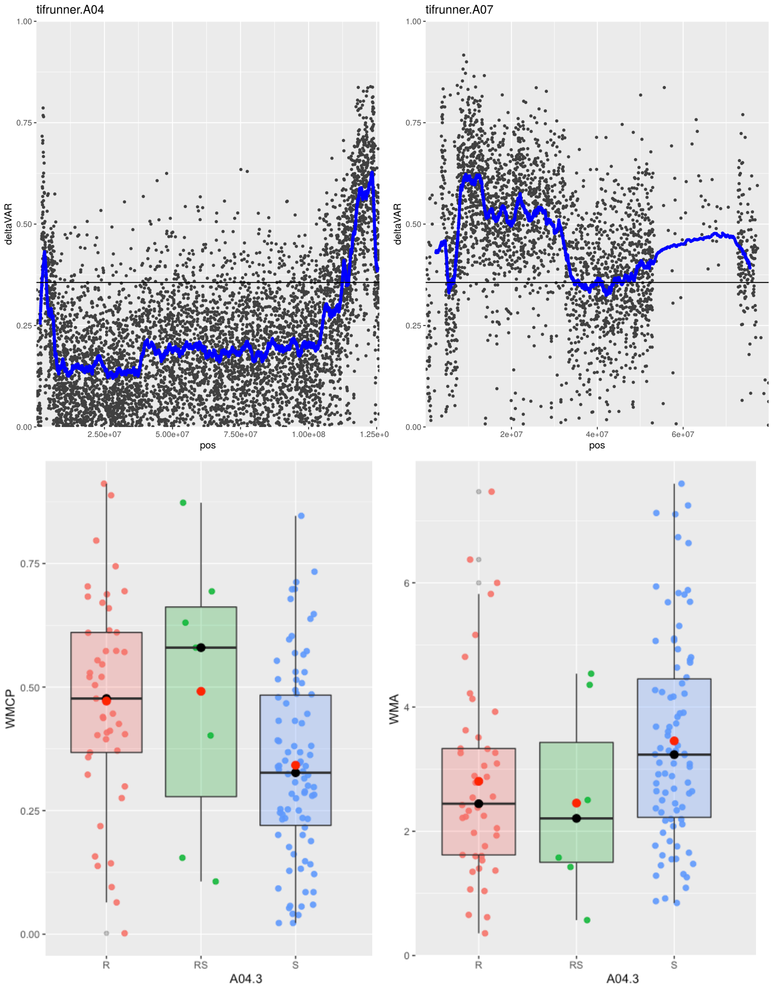


**Figure S2: Validation of additional minor QTLs identified.** Top – Additional potential QTLs identified by Khufu on A04 and A07. Bottom – Two measures of stem rot resistance in a population grouped by the putative beneficial (R) allele and susceptible (S) allele in the QTL region on A04 (red are individuals with the R allele, green are heterozygous, and blue have the S allele. WMCP represents the percentage of plants in a plot with a stem rot rating of 0 or 1 (no symptoms or 1 small lesion). WMA represents a disease score (0 – 10) of inoculated plants where 0 is no symptoms and 10 is a deceased plant. Scores are LSMeans from 3 years of data (2013, 2014, 2015) and 3 replicates per year. More information on the population and phenotypes available in Cui *et al*., 2019. For the WMCP trait, the QTL on A04 is significant using an ANOVA on ranks (F=6.95; p = 0.001) and a Kruskal Wallis non parametric test (p = 0.001). For the WMA trait, the QTl on A04 is significant using an ANOVA (F = 3.16; p = 0.046) and a Kruskal Wallis test (p = 0.036).

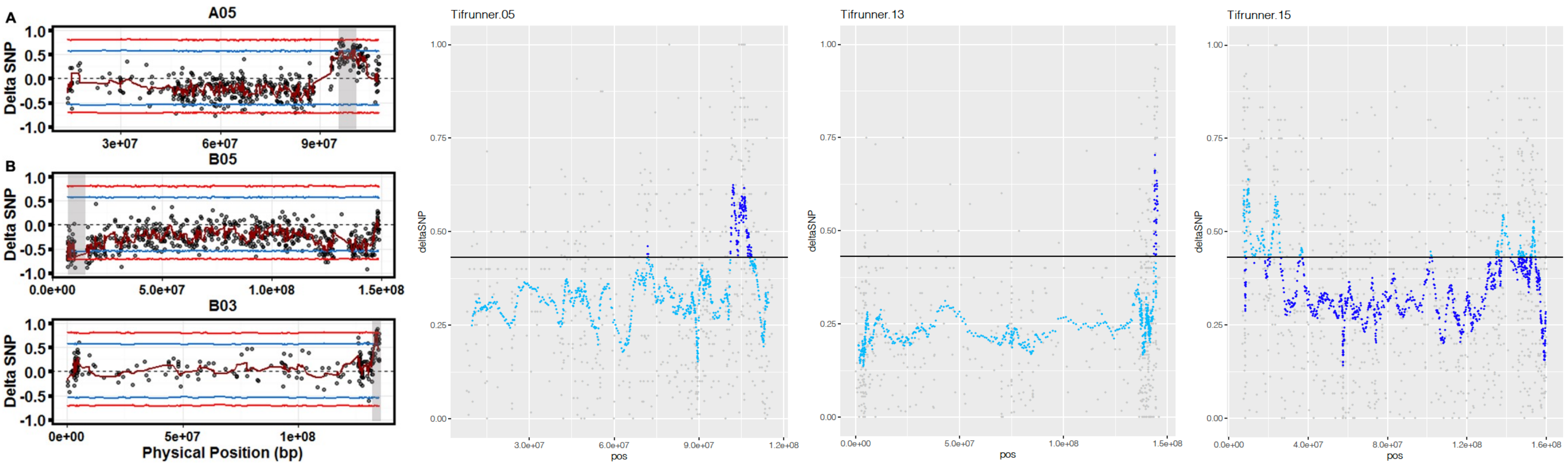


**Figure S3:** Scatter plots from Clevenger *et al*., 2017 with validated QTL on A05 (05), B05 (15), and B03 (13) using parental sequence. Khufu analysis straight from bulk sequence.

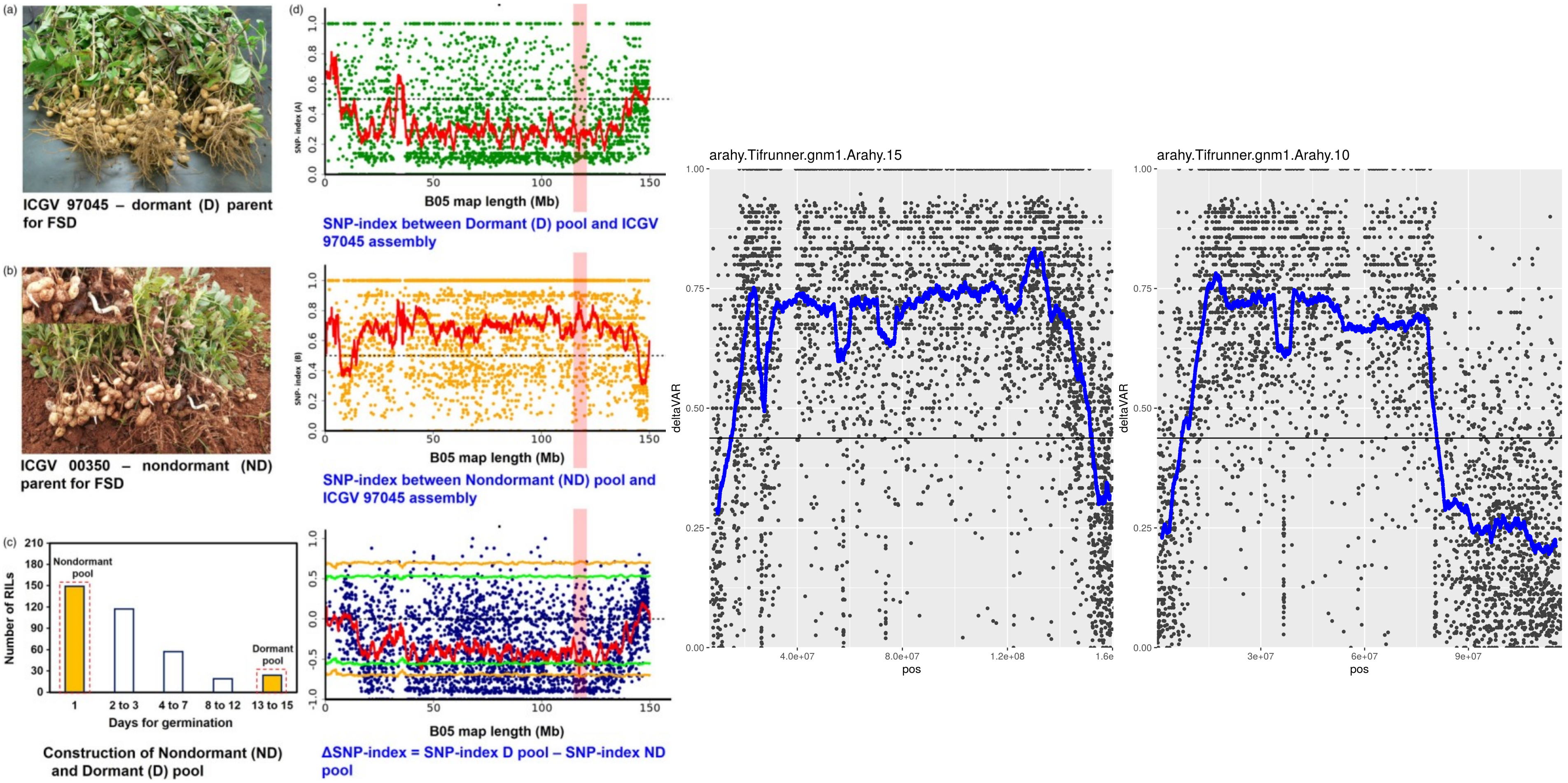


**Figure S4:** Figure from Pandey *et al*., 2020 showing the identification of a QTL controlling fresh seed dormancy on chromosome B05 (15) using parental sequence as a guide. The same data analyzed *de novo* with Khufu. Additional QTL were identified on chromosome 10 (shown) and 9 which were not reported in Pandey *et al*., 2020.

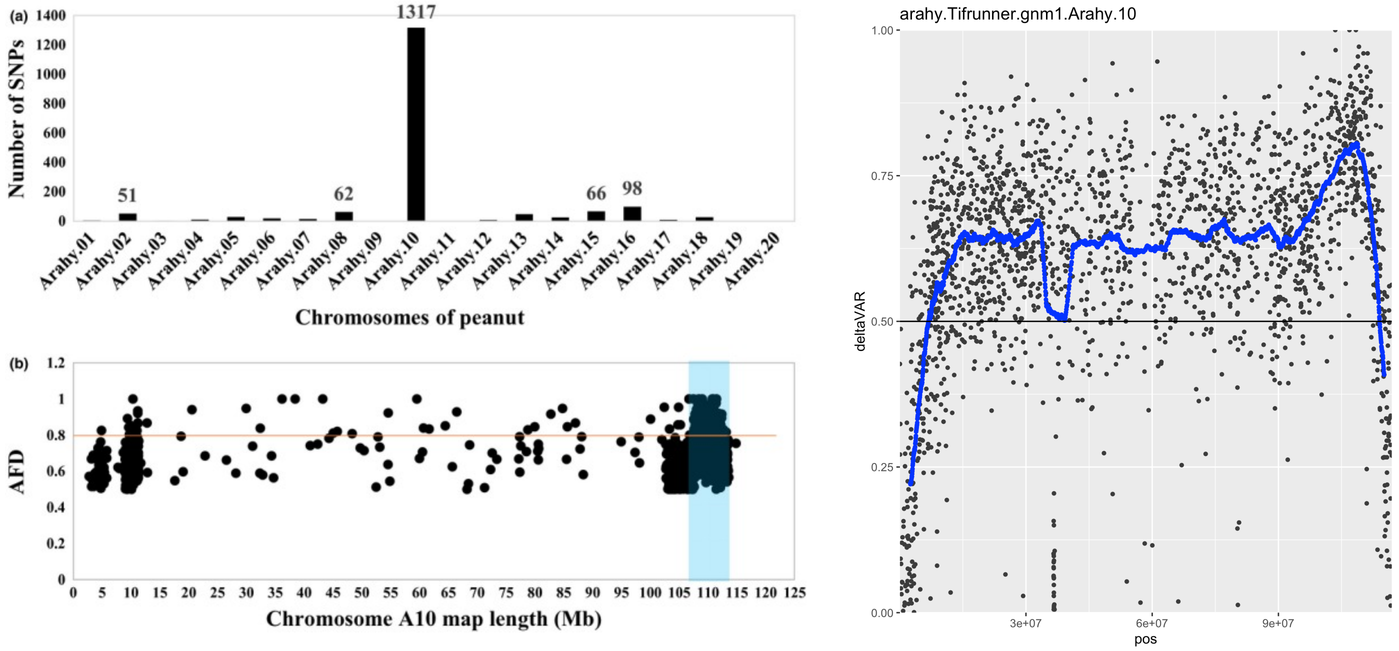


**Figure S5:** Left - Figure from Zhao *et al*., 2020 showing the identification of a QTL controlling purple testa color on chromosome A10 (15) using parental sequence. Right - The same data analyzed *de novo* with Khufu. The top of the peak indicates the region flanked by 108 Mb and 109 Mb contains the functional variation. Zhao and colleagues showed a myb transcription factor as the most likely candidate gene near 108.9 Mb.

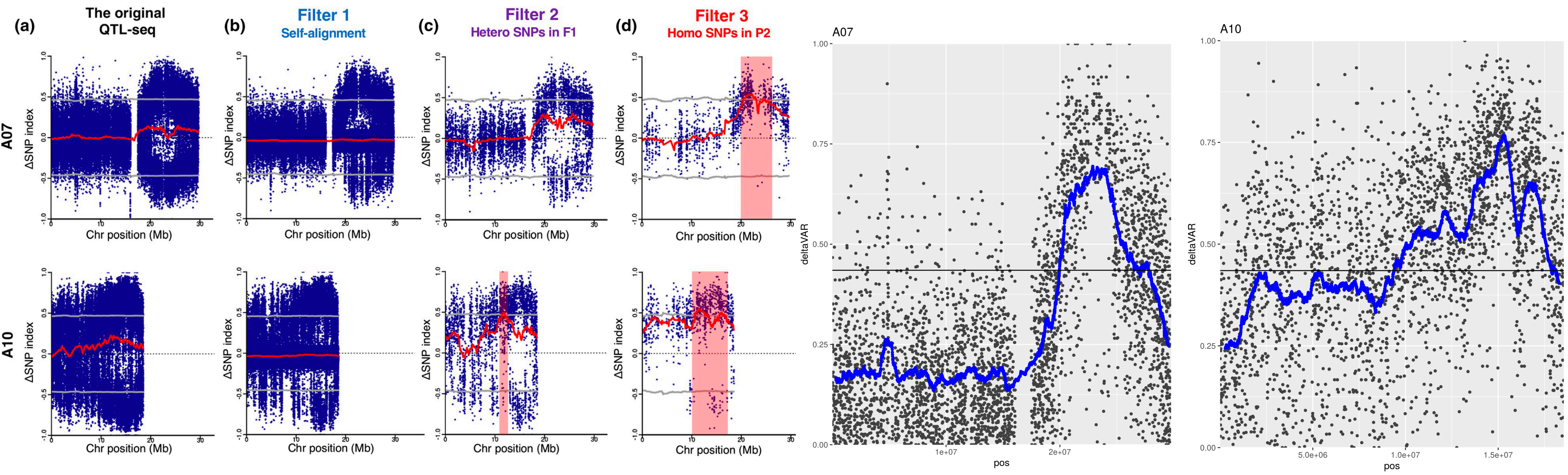


**Figure S6:** Left – Figure from Itoh *et al* (2019) showing their procedure for identifying QTL using sequence from parent 1 (filter 1), the F1 hybrid (filter 2), and parent 2 (filter 3). Significant QTL were identified on chromosomes A07 and A10. Right – Khufu identifies the same QTL *de novo* from the bulk sequence without needing additional sequencing.

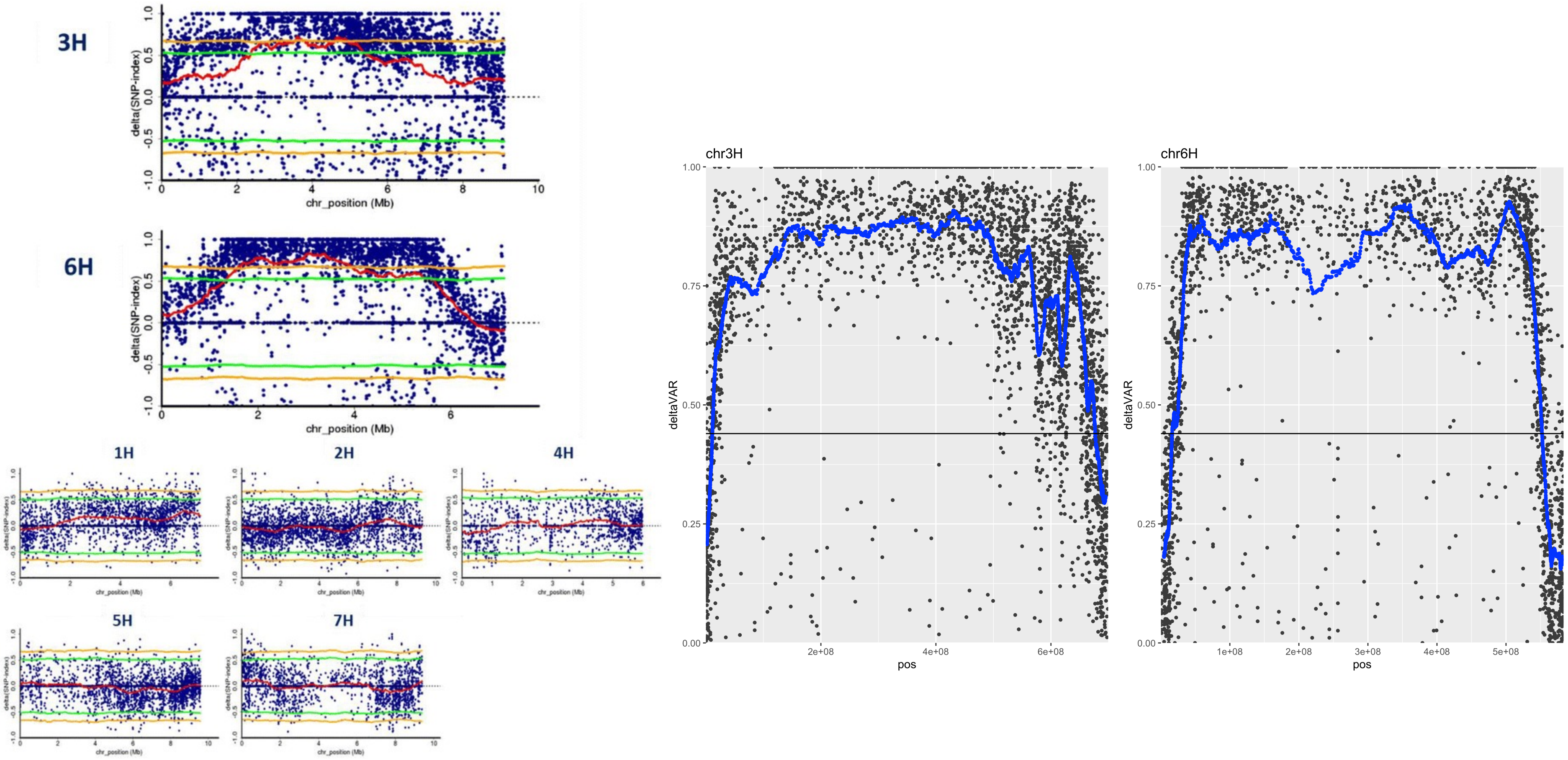


**Figure S7:** Left – Figure from Hisano *et al* (2018) showing QTL-seq for net blotch resistance. Significant QTL were identified on chromosomes 3H and 6H. Right – Khufu identifies the same QTL.

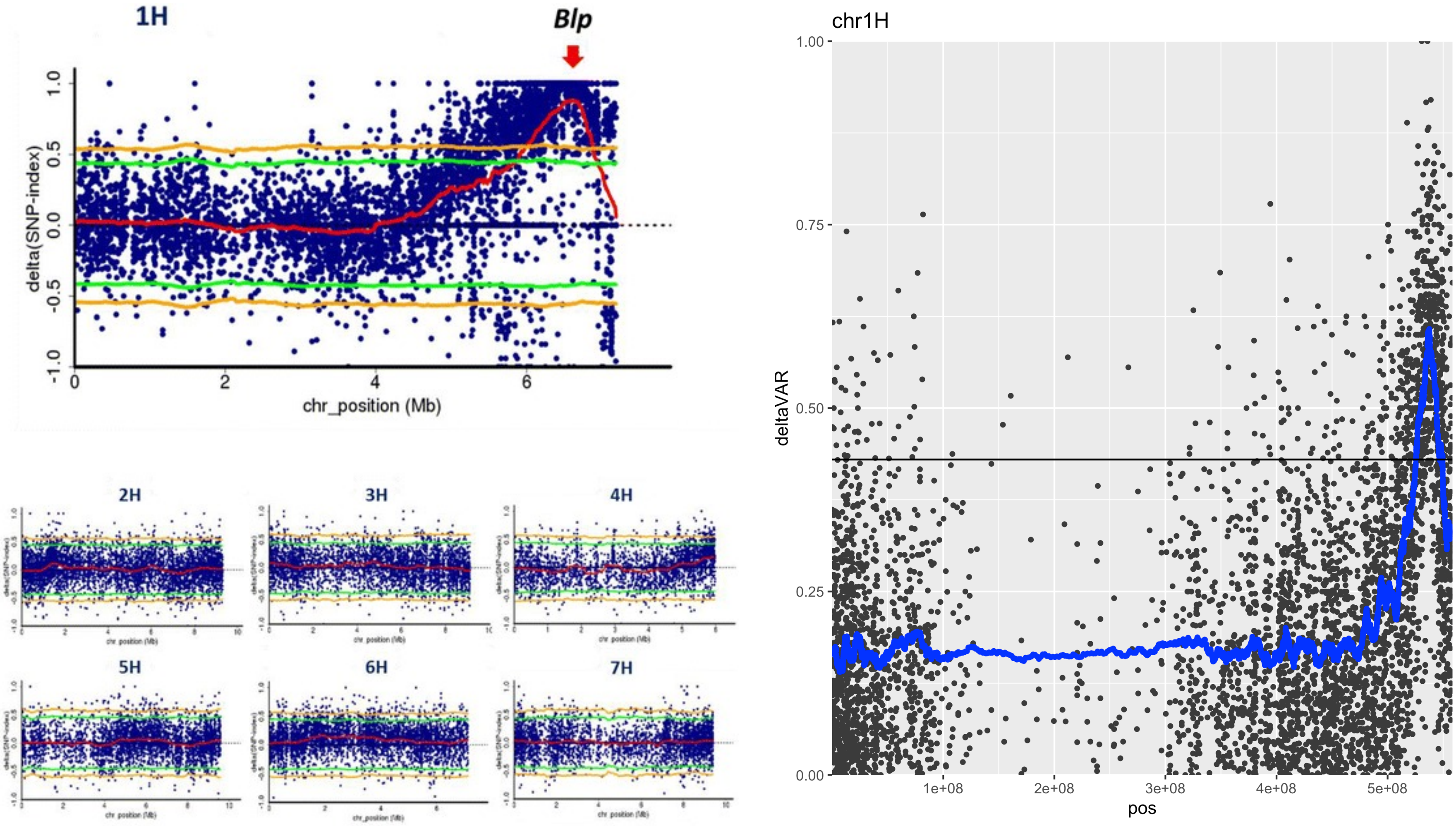


**Figure S8:** Left – Figure from Hisano *et al* (2018) showing QTL-seq for *blp*. Significant QTL were identified on chromosomes 3H and 6H. Right – Khufu identifies the same QTL. The position of the peak and top SNP is positioned in a candidate gene for *blp* identified in Long *et al*. (2019).

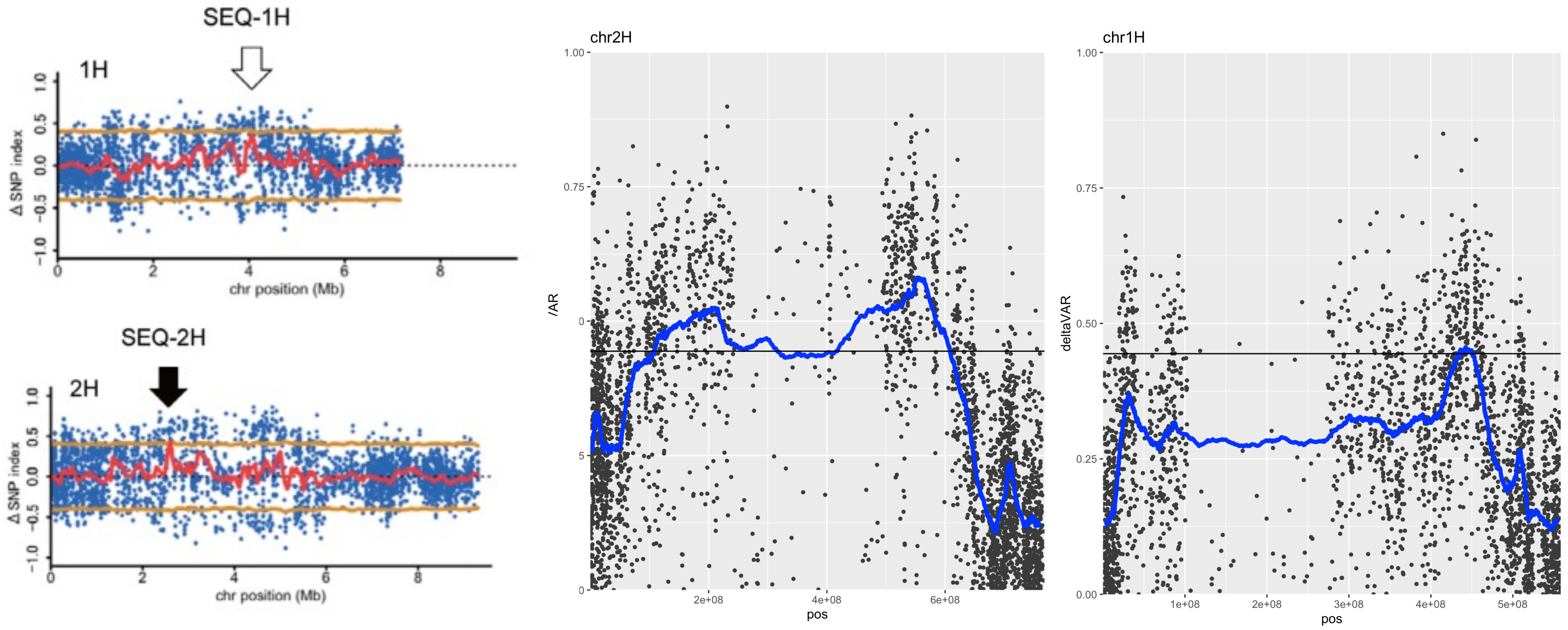


**Figure S9:** Left – Figure from Kodamo *et al* (2018) showing QTL-seq for salt tolerance in Barley. Significant QTL were identified on chromosomes 1H and 2H. Right – Khufu analysis.

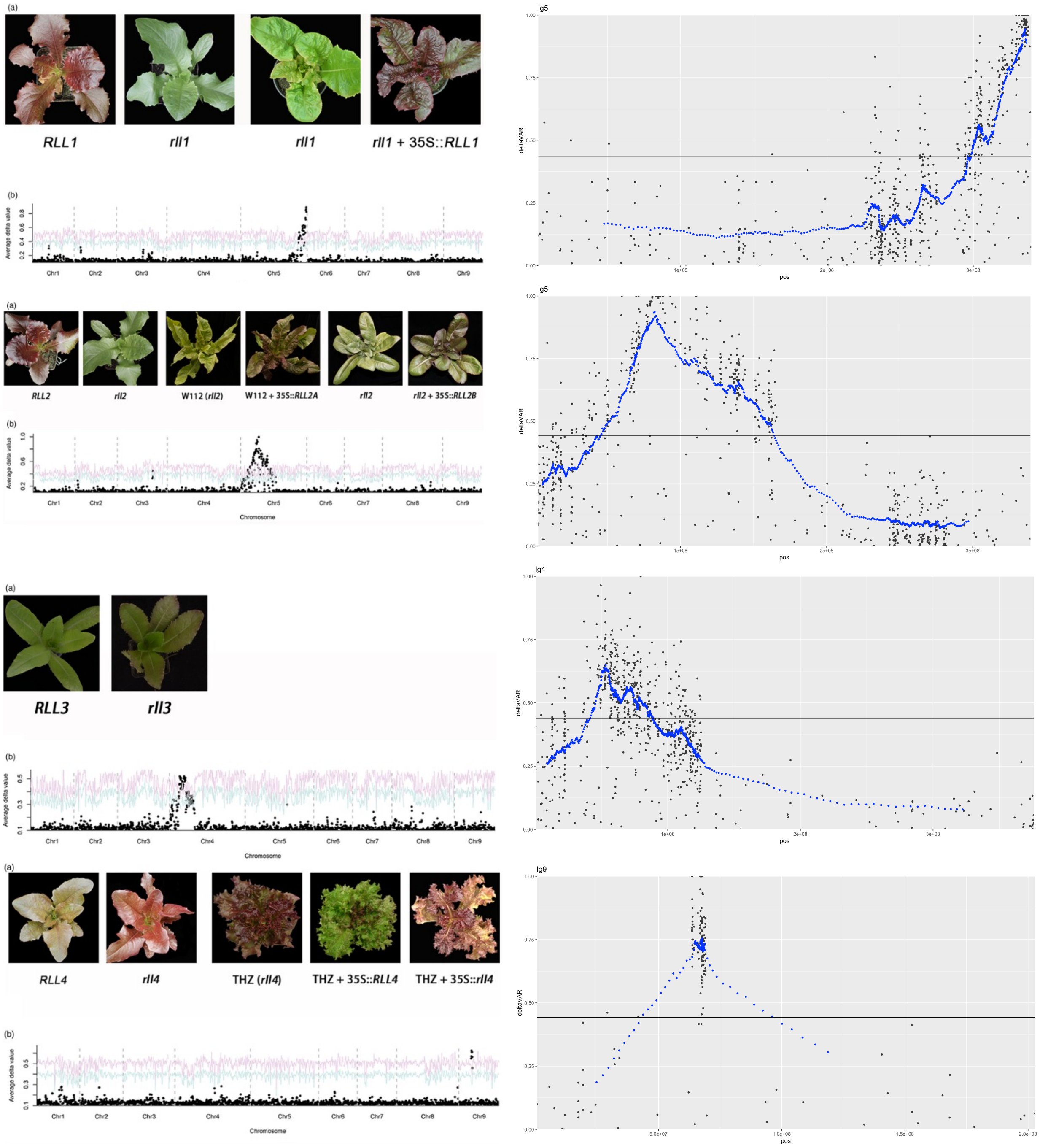


**Figure S10:** Left – Figure from Su *et al* (2020) showing QTL-seq using RNA sequencing to map 4 loci contributing to red color in lettuce. *RLL1* maps to the end of lg9; *RLL2* and *RLL3* map to lg5; *RLL4* maps to lg9. Right – Khufu analysis of the RNA-seq data.

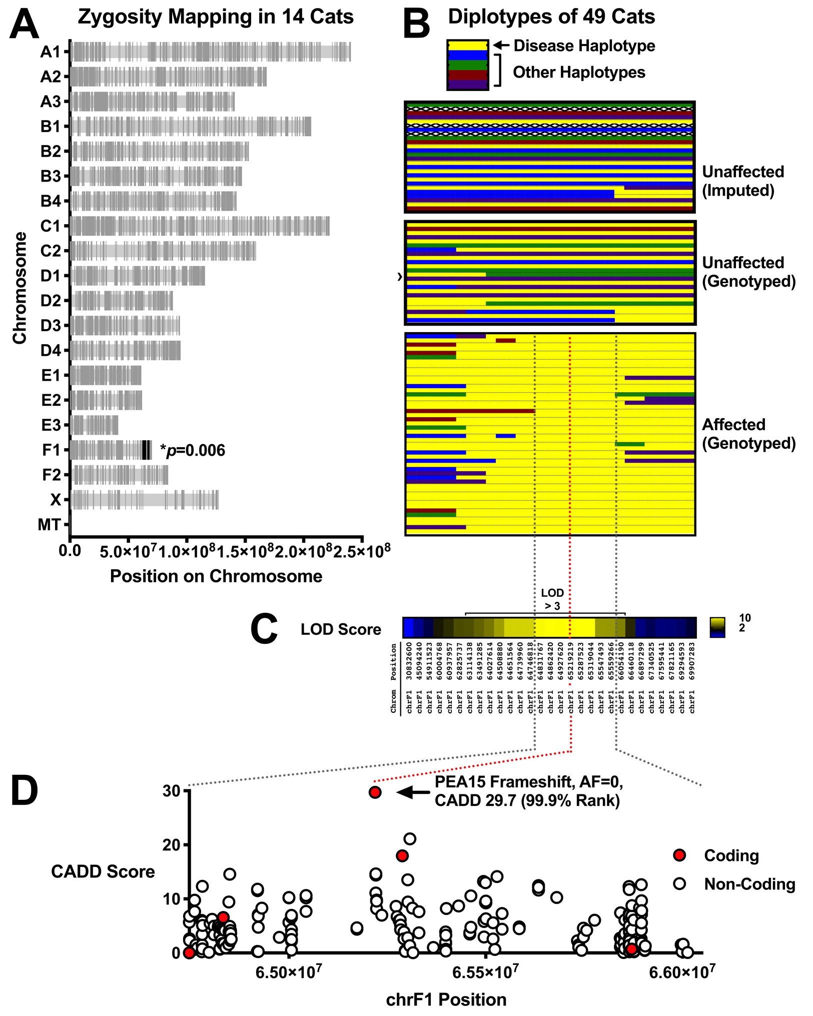

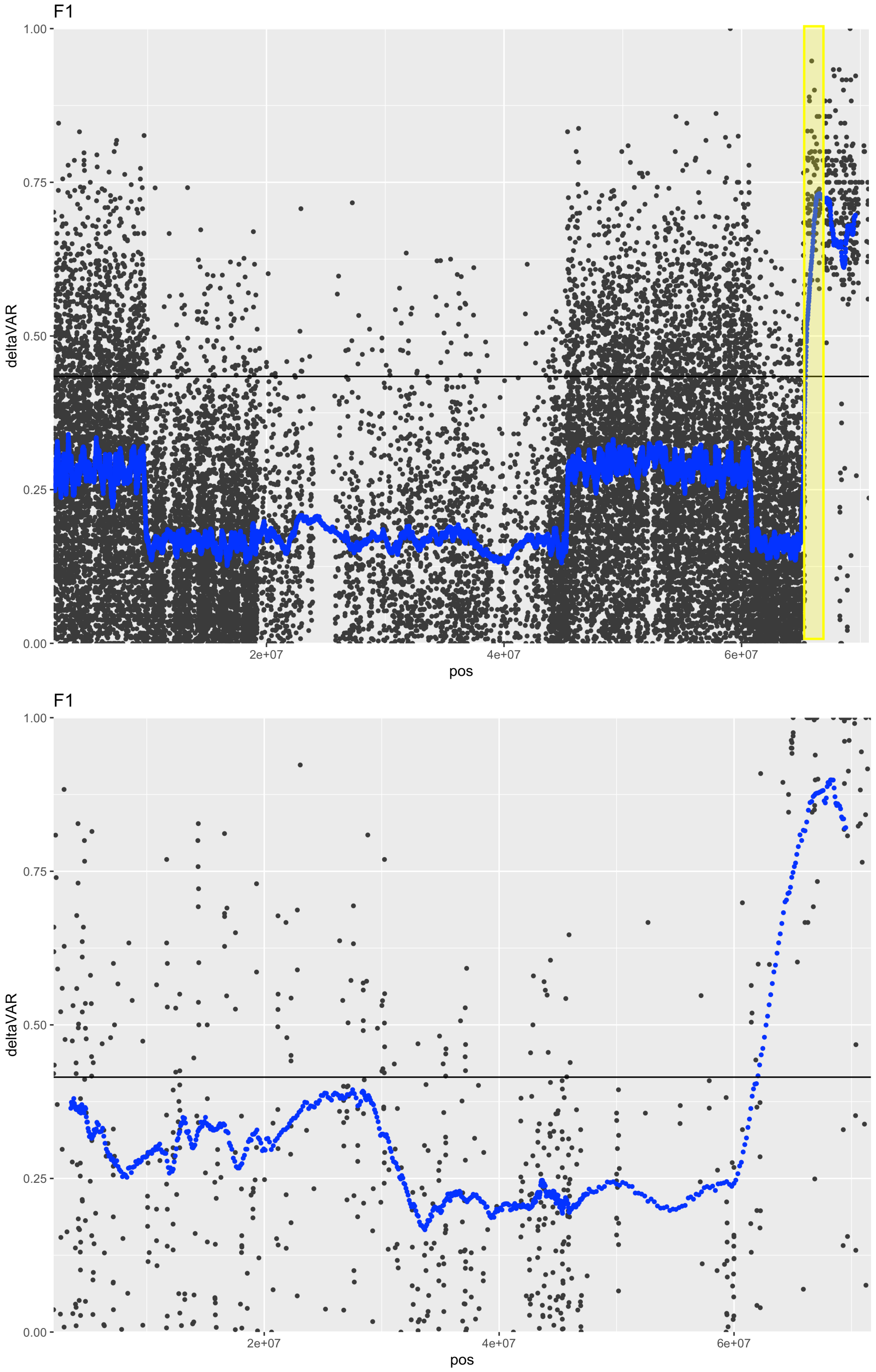


**Figure S11:** Left – Figure from Graff *et al* (2020) showing impaired cerebral cortical size in domestic cats. A mutation was identified in *PEA15* on chromosome F1. Right – Using just the cats with WGS from Graff *et al,* Khufu

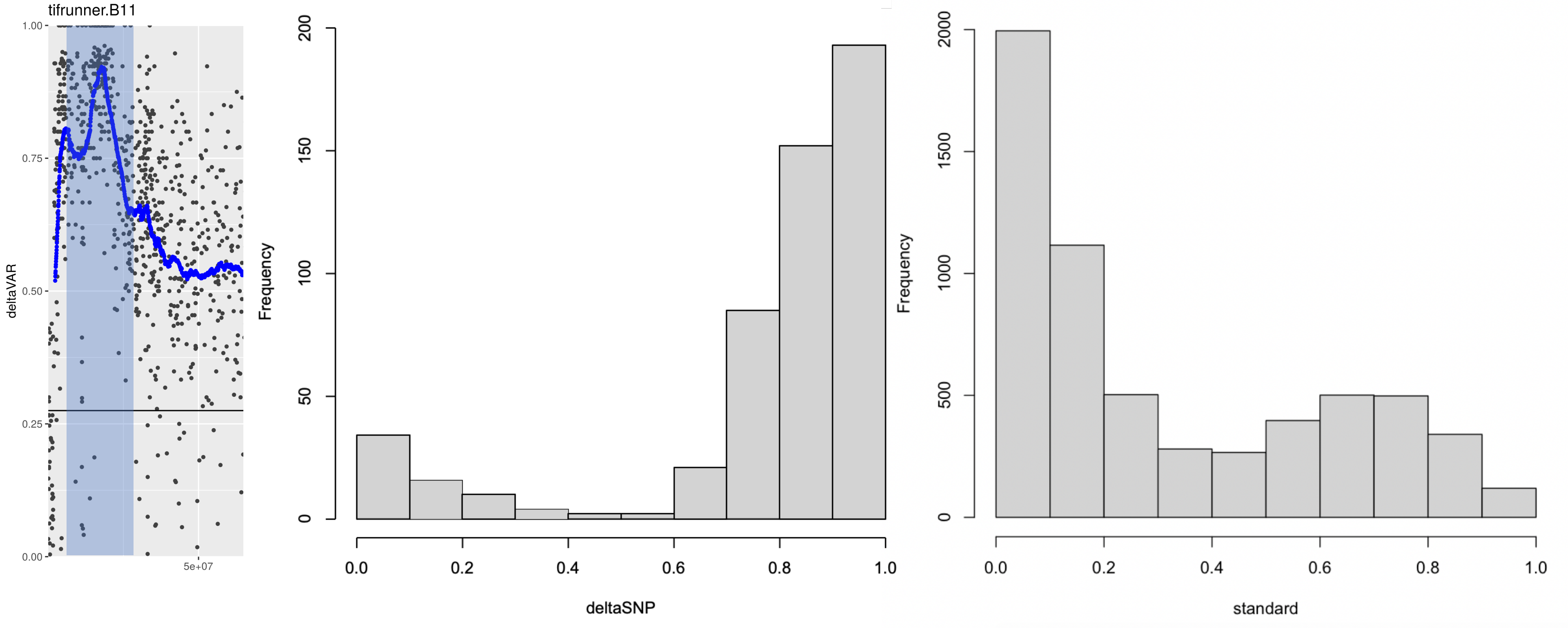


**Figure S12:** Distribution of delta SNP values from Khufu analysis (left) and a standard approach (right) within a peak on chromosome B01 (11) of peanut affecting blanching %.

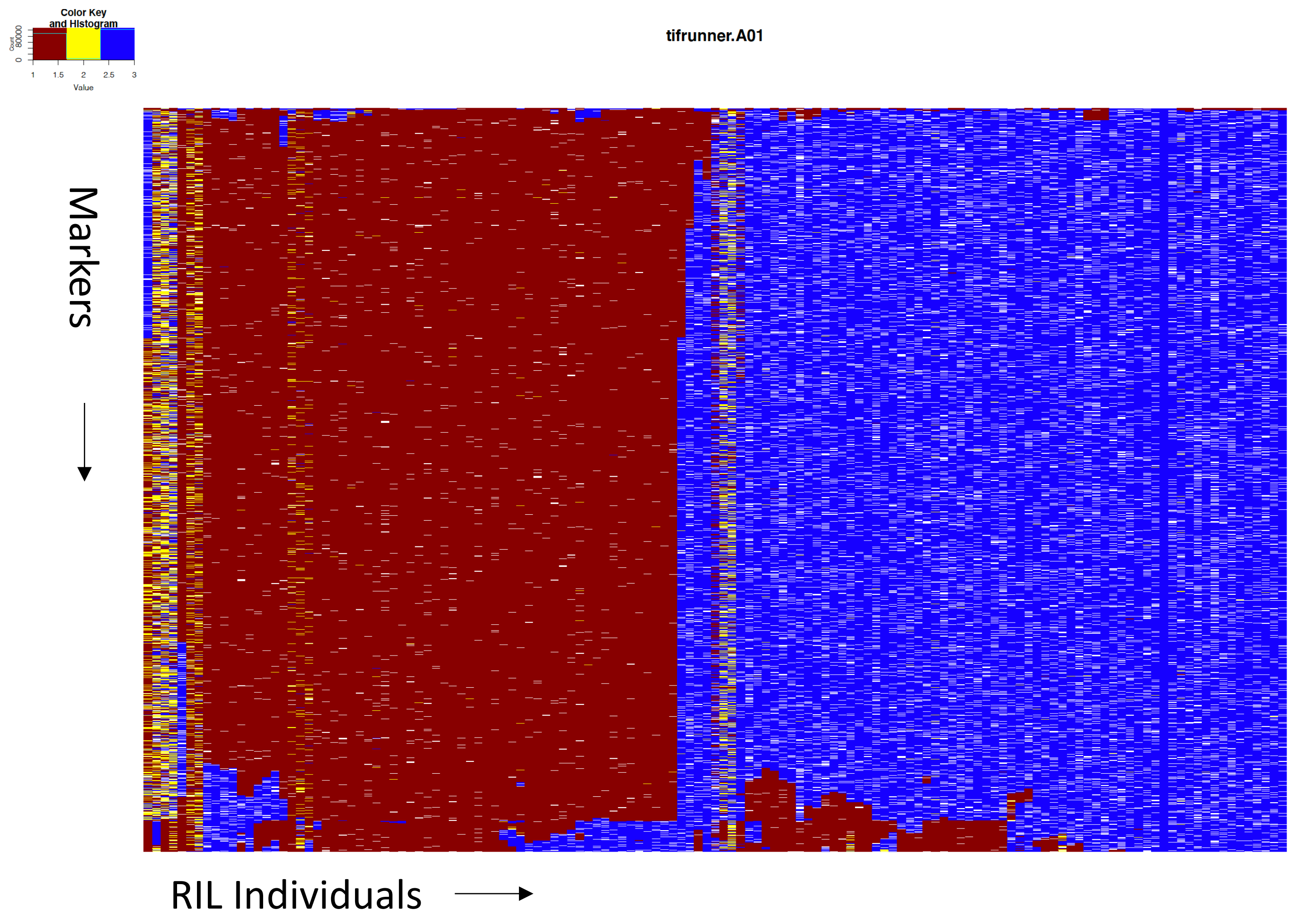


**Figure S13:** Representative physical map of 133 RIL individuals. Shown is chromosome A01. Red represents parent 1, yellow represents heterozygous, and blue represents parent 2. White is missing data. Individuals are clustered by similarity for visualization. Alleles are unfiltered out of the Khufu pipeline other than missing data <25%.

Table S1: Accuracy of allele calls using Khufu at different depths. Using SNPs identified from comparing reference genome sequences as a validation set, Khufu was used to call alleles using Illumina data at different coverages.

| Genotype | Correctly Called | Total Overlap | Accuracy | Khufu Called | Nucmer Identified | Depth |
|---|---|---|---|---|---|---|
| Tifruner | 34,443 | 34,594 | 99.56 | 287,809 | 74,885 | 3 |
| Shitouqi | 34,368 | 34,471 | 99.70 | 287,809 | 74,885 | 3 |
| Tifruner | 53,529 | 53,588 | 99.89 | 412,067 | 74,885 | 5 |
| Shitouqi | 53,273 | 53,388 | 99.78 | 412,067 | 74,885 | 5 |
| Tifruner | 69,230 | 69,235 | 99.99 | 399,010 | 74,885 | 10 |
| Shitouqi | 68,753 | 68,859 | 99.85 | 399,010 | 74,885 | 10 |
| Tifruner | 72,650 | 72,655 | 99.99 | 367,672 | 74,885 | 20 |
| Shitouqi | 71,909 | 71,992 | 99.88 | 367,672 | 74,885 | 20 |
| Tifruner | 73,190 | 73,193 | 100.00 | 360,454 | 74,885 | 37 |
| Shitouqi | 72,176 | 72,250 | 99.90 | 360,454 | 74,885 | 34 |

Nucmer identified 74,885 SNPs between te Shitouqi and Tifrunner genomes in regions that were not hard repeat masked. Khufu called a range of SNPs at different depths with an accuracy of allele calls >99% across all depths.

Recovery of SNPs at low coverage was 46% (3X)

**Table S2:** Polymorphic SNPs were called between Tfrunner and Shitouqi using 10X coverage Illumina sequence. To simulate "progeny", we downsampled the sequence to 0.5X and 1X coverage and called alleles base don the "parental" SNP sites.

| Depth | Polymorphic in Parents (10X) | Overlapped | Correctly called | Accuracy |
|---|---|---|---|---|
| 0.5 | 78,592 | 10,675 | 10,675 | 100% |
| 1 | 78,592 | 21,622 | 21,622 | 100% |

The number of overlaped SNPs was the number of sites at that coverage where an allele call was possible. The correct allele was called 100% of the time.
